# Bringing circuit theory into spatial occupancy models to assess landscape connectivity

**DOI:** 10.1101/2023.09.26.559320

**Authors:** Maëlis Kervellec, Thibaut Couturier, Sarah Bauduin, Delphine Chenesseau, Pierre Defos du Rau, Nolwenn Drouet-Hoguet, Christophe Duchamp, Julien Steinmetz, Jean-Michel Vandel, Olivier Gimenez

## Abstract

Connectivity shapes species distribution across fragmented landscapes. Assessing landscape resistance to dispersal is challenging because dispersal events are rare and difficult to detect especially for elusive species. To address these issues, spatial occupancy models have been developed to integrate the resistance surface concept of landscape ecology and model patch occupancy dynamics through colonization and extinction while accounting for imperfect species detection. However, the most recent approach is based on least-cost path distances which assume that individuals disperse along the optimal route. Here, we develop a new spatial occupancy model that incorporates commute distances derived from circuit theory to model dispersal across sites. Our approach allows for explicit estimation of landscape connectivity and direct measure of uncertainty from detection/non-detection data. To illustrate our approach, we study the recolonisation of two carnivores in France, and quantify the degree to which rivers facilitate Eurasian otter (*Lutra lutra*) dispersal and highways impede Eurasian lynx (*Lynx lynx*) recolonisation. Overall, spatial occupancy models provide a flexible framework to acccommodate any distance metric designed to align with species dispersal ecology.

**Open Research Statement:** Data and code used in this research are available on Zenodo at https://zenodo.org/record/8376577

## 1. Introduction

The way a landscape is shaped structures ecological processes at different spatial and temporal scales, from within individual home-range movement to species range (Crooks et Sanjayan 2006). Understanding landscape connectivity has motivated the development and use of different approaches. Some authors have characterised landscape structure, while others have quantified how spatial distribution of patches affects colonisation and extinction (Moilanen et Hanski 2001; Tischendorf et Fahrig 2001). By linking landscape ecology and metapopulation theory (Howell et al. 2018), ecologists may gain valuable insights into how disruptions in connectivity, e.g. habitat fragmentation, impact movement (Hilty, Lidicker, et Merenlender 2006), with important implications for species conservation and management.

In metapopulation theory, the colonization of new habitat patches is primarily accomplished through dispersal events (Hanski et Gilpin 1991). However, these events are rare which makes them hard to detect. To address this issue, dynamic occupancy models have been developed to assess species distribution patterns while accounting for imperfect detection by using detection and non-detection data, and to explicitly estimate the probabilities of colonization and extinction (MacKenzie et al. 2017). While these probabilities can be function of patch characteristics, their spatial arrangement in the landscape is commonly ignored. Chandler et al. (2015) extended the standard dynamic occupancy model by considering the Euclidean distance between patches in the colonisation process. This so-called spatial occupancy model allows for a more realistic representation of how species disperse, by acknowledging that a site close to occupied sites is more likely to be colonised. However, this model does not account for resistance of the matrix in between patch movements. To cope with this issue, Howell et al. (2018) replaced the Euclidean distance by the least-cost path (LCP) distance which is often used in landscape ecology (Zeller, McGarigal, et Whiteley 2012; Coulon et al. 2015). This model allows for explicit estimation of cost values and connectivity. An important assumption of the LCP approach is that all individuals are omniscient and will follow the optimal route. However, in situations where dispersers do not have previous knowledge of the landscape, the omniscience assumption does not hold (Diniz et al. 2020).

To relax this assumption, circuit theory has been introduced as a conceptual framework that applies principles from electrical circuit theory to model the movement of individuals across the landscape, like electrons in a circuit (McRae et al. 2008). In circuit theory, the landscape is considered as a network of interconnected nodes linked by resistors that form the resistance surface. By assigning resistance values to each landscape feature, circuit theory quantifies the potential flow of individuals between patches. This formulation implies that an individual moving through the landscape ignores the optimal route between patches and makes decisions at each step. In circuit theory, commute distance is used to compute inter-patches distances. This distance is defined as the expected number of steps a random walker would take to move between patches and return. The commute distance has the advantage to account for path redundancy, in other words the commute distance between patches decreases when there are multiple ways to join these patches (McRae et al. 2008). Despite many applications of circuit theory in landscape ecology, the commute distance is yet to be brought into the occupancy framework.

In this paper, we integrate the commute distance in the colonisation process of dynamic occupancy models. We adopt a hierarchical representation of occupancy models (MacKenzie et al. 2017) which we implement in the Bayesian framework using Markov chain Monte Carlo methods in the R package NIMBLE (de Valpine et al. 2017). To illustrate our approach, we use two case studies on carnivores recolonisation in France: the Eurasian otter (*Lutra lutra*) in the Massif Central and the Eurasian lynx (*Lynx lynx*) in the Jura mountains. Using our model and long-term detection/non-detection data, we aim to quantify the degree to which rivers facilitate otter dispersal and highways impede lynx recolonisation.

## 2. Material and Methods

### 2.1 Spatial dynamic occupancy model with commute distance

Dynamic occupancy models rely on detections and non-detections, denoted *y_i,j,t_*, of a species at site *i* on survey *j* during season *t*. Because the true presence-absence *z_i,t_* of a species at site *i* during season *t* is only partially observed, we accommodate for imperfect species detection with probability p*_i,j,t_*, which we estimate from repeated surveys:

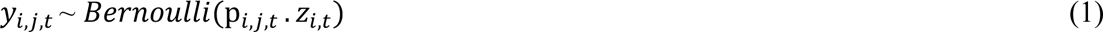

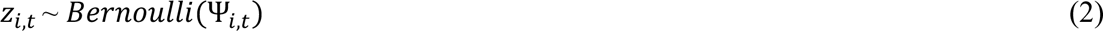

where Ψ*_i,t_* is the occupancy probability at site *i* during season *t*. Probabilities of colonisation and extinction are denoted *γ* and *ε*. A site is occupied in a given season if it was unoccupied at the previous season and is colonised, or if it was occupied in the previous season and did not get extinct:

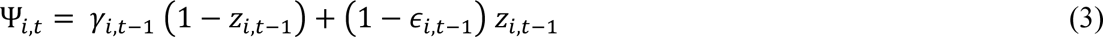

Parameter *γ*_*i,t*–1_, the probability of a site *i* to be colonised between season *t* − 1 and *t*, is defined as the complementary of the product of the probabilities of not being colonised over all occupied sites *m* in the landscape, where M is the total number of sites:

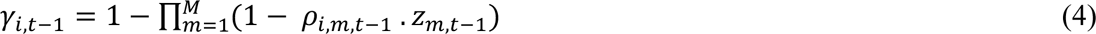

The probability *ρ*_*i,m,t*–1_ that site *i* is colonised by site *m* between season *t* − 1 and *t* is called pairwise colonisation probability (Chandler et al. 2015), and is modelled wih a gaussian kernel. This approach is the standard way to have a probability that decreases with increasing distance between sites *d_i,m_*:

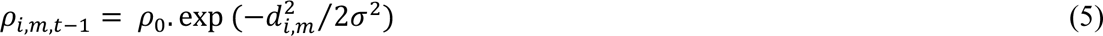

where *ρ*_0_ is the baseline pairwise colonisation probability when *d_i,m_* = 0 and *σ* is the scale parameter that describes the colonisation distance.

The originality of our model lies in the use of a commute distance *d_i,m_* instead of a Euclidean distance (Chandler et al. 2015) or a LCP distance (Howell et al. 2018). In circuit theory, distances are computed on a resistance surface, which is obtained using a landscape raster (McRae et al. 2008). From this raster, we define a graph where each pixel is a node connected to the eight adjacent nodes with resistors. The resistance or cost between two adjacent nodes *x* and *x’* is function of the value of the landscape variable *c*_1_ at these nodes and an estimated resistance parameter *α* ∈ ℝ:

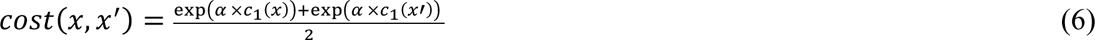

When *α* > 0, the landscape is assumed to impede movement, whereas the more alpha is negative, the more the landscape facilitates movement. Overall when *α* = 0, the resistance between all adajacent nodes is the same and the landscape structure has no influence on movement. Given a source and a destination in the graph, the commute distance is the average number of steps needed to join the two nodes and back during a random walk. The commute distance is computed as two times the number of edges in the graph times the effective resistance between the two nodes (Chandra et al. 1997). To compute commute distances, we used the commuteDistance() function of the gdistance R package (van Etten 2012). All analyses were conducted in R (version 4.2.2, R Core Team 2022). We implemented the spatial occupancy model with commute distances in the Bayesian framework using NIMBLE (version 0.13.1, de Valpine et al. 2017). NIMBLE allows to call any R functions like commuteDistance() from within a model, which made the implementation of our model possible. We provide the NIMBLE code for dynamic occupancy models, models accommodating Euclidean distances, least-cost path distances and our model with commute distances (Appendix S2).

### 2.2 Applications

#### 2.2.1 European otter population in the Massif Central

We focus on the Eurasian otter population distributed in the Massif Central in France (Figures 1 a,b). We used detection/non-detection data from 158 sites in the Massif Central that were previously analysed by Couturier et al. (2023). The sites are 300m transects along rivers where the centre is placed at potential marking locations (e.g. bridges, water mills, confluences). The monitoring is composed of two winter seasons, with the first one lasting two consecutive winters, from November to April in 2003-2004 and in 2004-2005 (season 1) and from October 2011 to April 2012 (season 2). Otters in the Massif Central are genetically close (Pigneur et al. 2019), suggesting that the new colonized sites are likely to be from individuals already present in the massif. Otter recolonisation is a slow process, making the assumption of closure within a season most likely valid. Each site is visited between 1 and 4 times in the first season and between 2 and 3 times in the second season.

**Figure 1.**
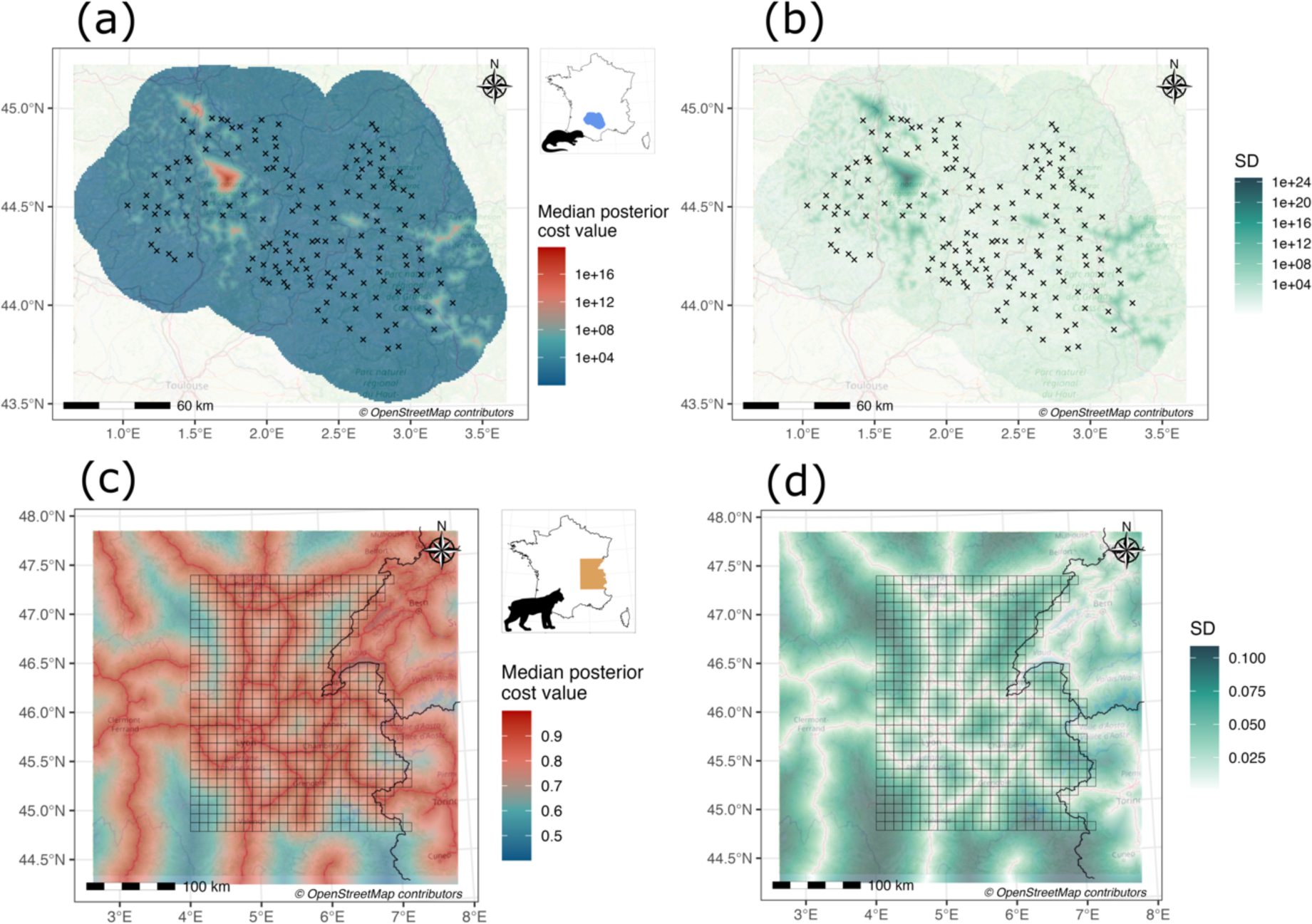
- Distribution of sampling sites for the two case studies in France. On the top panels, black crosses symbolize the center of 300m transect monitoring otter spaint in the Massif Central, background raster represents the median posterior cost value estimated from the distance to rivers (a) and corresponding standard deviation (b). On the bottom panels, 100km^2^ squares delineate sampling sites of lynx presence in the Jura mountains, background raster represents the median posterior cost value estimated from the distance to highways (c) and corresponding standard deviation (d).

We model initial occupancy probability as a function of elevation (BD ALTI^®^ Version 2.0, 01/2023) because otters are assumed to colonize from mountains (Appendix S1: Figure S1 a). Detection probability is assumed to be season-specific (Couturier et al. 2023). Because otters mainly use rivers to colonise new habitats, we compute the distance to the closest river (BD TOPO^®^ Version 3.3, 12/2022) to create the resistance surface with a resolution of 0.04km^2^. This surface encompasses all sampling sites with a 30km buffer around them to account for all possible alternative routes between sites.

We run three chains each of length 10,000 and we discard the first 1,000 iterations as burn in. We use non-informative priors for all parameters except for the resistance parameter (Appendix S2) for which we use a uniform distribution between 1 and 5, because otters are semi-aquatic and do not go far away from rivers, and therefore the larger distances to rivers is assumed to strongly impede their movements (Couturier et al. 2023). Given that the landscape covariate is reduced and that the resistance scale is exponential, we set the upper bound to 5 to ensure that the commute distance between all sites is not too small.

#### 2.2.2 Eurasian lynx population in the Jura mountains

We focus on the lynx population present in the French Jura mountains (Figures 1 c,d). We use a 100 km^2^ grid with 644 cells in the French Jura where sampling occured at least once over the study period. Because lynx recolonisation is a slow process, we define 3 seasons at 10-year intervals in 1999-2000, 2009-2010 and 2019-2020. We consider months from November to April as replicated surveys. A detection occurs at a sampling occasion if a sign of lynx presence (e.g., tracks, hair, camera trap picture) is found in a cell of the grid. Then the detection is allocated to the site defined as the center of the grid cell. The monitoring is carried out by a network of observers trained by the French Office for Biodiversity. We assume that the detection probability vary according to sampling effort calculated as the number of observers present in a cell in a given season and survey (Louvrier et al. 2018). The lynx is an elusive species living in forested areas where its main preys, ungulate species, occur (Molinari-Jobin et al. 2007). Therefore we assume that initial occupancy probability is a function of forest cover which we retrieve from Corine land cover data (European Environment Agency, 2000) (Appendix S1: Figures S1 b). We consider the distance from the center of the pixel to the nearest highway computed from Open Street Map data (OpenStreetMap contributors 2023) as the resistance surface with a resolution of 6.25km^2^. This surface encompasses the sampling grid with a 50km buffer.

We run three chains each of length 10,000 and we discard the first 5,000 iterations as burn in. We use non-informative priors for all parameters except for the resistance parameter (Appendix S2), for which we use an uniform distribution between −5 and 0 because short distance to highways is assumed to limit lynx dispersal (Kramer-Schadt et al. 2004). Given that the landscape covariate is reduced and that the resistance scale is exponential, we set the lower bound to −5 to avoid extreme commute distance values.

## 3. Results

We detect otter presence at 122 out of the 158 sites over the 2 seasons and we detect lynx presence at 128 out of the 644 sites surveyed over the 3 seasons (Appendix S1: Figure S2). Hereafter, we provide the median of parameter posterior distribution along with 95% credible intervals. The two models converge according to visual inspection of trace plots and Gelman– Rubin statistic (Rhat ≤ 1.02; Appendix S3 & S4 ; Brooks and Gelman 1998). Otter detection probability is high over the whole study area, respectively 0.58 [0.49; 0.66] and 0.80 [0.74; 0.85] in the first and second sampling season (Appendix S1: Figure S3 a). Otter initial occupancy probability increases with increasing elevation from 0.29 [0.17; 0.43] at 75m to 0.89 [0.67; 0.98] at 1280m (Appendix S1: Figure S3 b). Lynx detection probability slightly increases with increasing sampling effort from 0.27 [0.23; 0.32] to 0.36 [0.23; 0.52] (Appendix S1: Figure S3 c). Initial lynx occupancy probability sharply increases with increasing forest cover, and ranges from 0.01 [0.00; 0.02] without forest cover to 0.59 [0.43; 0.73] at 81% of forest cover (Appendix S1: Figure S3 d).

Both species have low extinction probability, 0.05 [0.01; 0.13] and 0.32 [0.21; 0.42] respectively for otter and lynx. Baseline pairwise colonisation probability for otter and lynx is contrasted and respectively equal to 0.02 [0.01; 0.06] and 0.94 [0.69; 1] for a given occupied site (Appendix S1: Figures S4 a,b). Recolonisation is clearly linked to habitat structure, with the estimated resistance parameter being 3.9 [1.16; 4.96] and −0.18 [−0.10; −0.29] for otter and lynx respectively. When inspecting the estimated cost surfaces, rivers facilitate otter dispersal, while highways are barriers for lynx dispersal (Figure 1). The further the site is from the rivers or highways, the greater is the cost uncertainty (Figures 1 c,d).

We also map the occupancy probability for otter and lynx (Figure 2) along with associated uncertainty (posterior standard deviation). In both case studies, the area occupied increases over the study period. In the Massif Central, almost all sites are occupied by otters in 2011, and only a few sites in the western part of the study area remain unoccupied (Figure 2 b). In the Jura mountains, lynx occupancy is bounded by the highways. As the distance from the core of the population increases, uncertainty decreases (Figure 2 j).

**Figure 2.**
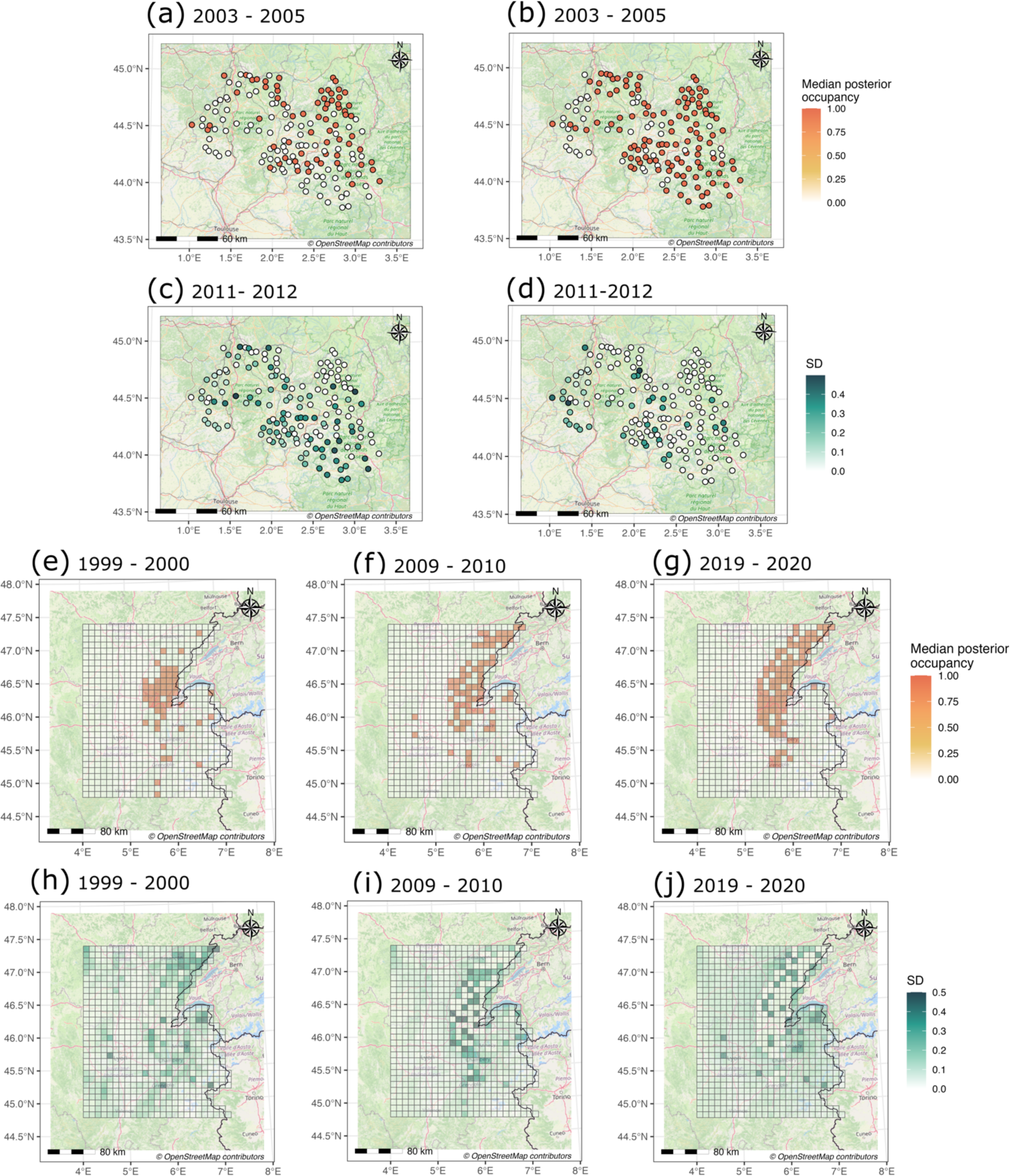
– Maps of estimated posterior occupancy probabilities for otter in the Massif Central at the first monitoring season in 2003 - 2005 (a) and second monitoring season in 2011-2012 (b) and for the lynx in the Jura mountains at the first monitoring season from November 1999 to April 2000 (e), the second monitoring season from November 2009 to April 2010 (f) and the third monitoring season from November 2019 to April 2020 (g) with respective standard deviation (c, d, h, i & j).

## 4. Discussion

Dynamic occupancy models offer a comprehensive understanding of species distribution over time while accounting for imperfect detection (MacKenzie et al. 2017). We extend existing models by integrating commute distance from circuit theory, therefore relaxing the assumption of omniscience and accounting for multiple dispersal routes (McRae et al. 2008). Our spatial occupancy model provides valuable insights into potential barriers to species dispersal and the identification of ecological corridors. Unlike traditional approaches that rely on expert opinion to define resistance costs (Compton et al. 2007; Zeller, McGarigal, et Whiteley 2012), our spatial occupancy model estimates resistance and connectivity based on empirical data and provides a direct measure of uncertainty.

In the two case studies, we find differences in strength and sign of landscape resistance on dispersal. The estimated resistance is positive for the otter regarding the distance to the rivers, which is consistent with the semi-aquatic nature of this species (Van Looy et al. 2014). The presence of fewer rivers in the Lot department results in slower colonization of the western sites. In contrast, the resistance parameter is positive for the lynx population regarding the distance to highways, and population range in the last season is clearly shaped by these linear features (Figure 2 j). However, we do not include other landscape covariates (e.g. forest cover, elevation) in the colonisation model and the habitat in the western side of the highways does not seem favourable for lynx establishement. Other landscape covariates correlated with highways distribution might also limit lynx dispersal to the west. While our model can be used to inform strategies for preserving or enhancing landscape connectivity, multiplying approaches and validating outputs from multiple data sources is also needed (Zeller et al. 2018; Riordan-Short, Pither, et Pither 2023). For instance, our approach focuses on dispersal in the sense of metapopulation theory (Hanski et Gilpin 1991), and does not account for gene flow or road mortality (Bauduin et al. 2021). If we were to evaluate the long-term viability of lynx and otter populations, these issues should be addressed and would require genetic and mortality data. Spatial occupancy models were originally developed for metapopulations (Chandler et al. 2015). Here, we apply these models to a single population in a continuous landscape. The sites do not represent a population or a patch anymore but a unit in a landscape, e.g. a pixel in a grid. This occupancy surface representation allows us to map precisely the distribution of the population within a season. For territorial species like carnivores, the occupancy surface resolution is defined according to home-range size (MacKenzie et al. 2017). However, animal distribution and animal dispersal are two processes that occur at different spatial scales and are affected by diverse variables, which explains why we use two distinct surfaces: the occupancy surface and the resistance surface. To model the movement process between seasons, the resolution of the resistance surface has to be the most refined possible to provide a realistic representation of potential dispersal routes (Zeller, McGarigal, et Whiteley 2012; Howell et al. 2018). Moreover, two key points must be taken into account when defining the resistance and occupancy surfaces. First, the resistance surface should include a buffer large enough to cover all possible paths between sites (Howell et al. 2018). Second, the occupancy surface should contain all sites that can be source of individuals, even those which are not monitored. In spatial occupancy models, dispersal is modelled as pairwise colonisation probability, meaning that the population is closed, and sites can be colonised only by individuals from sites present in the model (Chandler et al. 2015).

In the context of landscape connectivity, both LCP distance and commute distance offer distinct advantages and limitations. The LCP distance assumes that individuals have a perfect knowledge of the landscape and will follow the optimal route, making it a valuable metric for understanding the most efficient dispersal paths (Adriaensen et al. 2003). On the other hand, commute distance links animal movement via random walk where decision is made step by step. By accounting for multiple dispersal paths, the commute distance approach highlights pinch points in the landscape to ensure connectivity (McRae et al. 2008; Coulon et al. 2015). The model implementation is facilitated by NIMBLE (de Valpine et al. 2017) which allows to call R functions from within the model, making it possible to compute Euclidean distance, LCP distance and commute distance (Appendix S2). In reality, accurate modelling of dispersal path likely falls in the continuum between perfect knowledge (LCP) and complete ignorance (commute distance) of the landscape structure. The randomized shortest path solves this trade-off at the cost of an additional parameter, *θ* ∈ ℝ^+^, to estimate. When *θ* = ∞, this distance is equivalent to LCP distance and when *θ* = 0, the distance is equivalent to commute distance (Panzacchi et al. 2016). We note that any distances could be included in the model, like spatial absorbing Markov chains (Fletcher et al. 2023) or individual-based dispersal models (Coulon et al. 2015).

The flexibility of our framework makes it appealing to introduce additional complexity. We recommend selecting the dispersal model and landscape variables that best align with the ecology of the focal species and the research question (McClure, Hansen, et Inman 2016; Diniz et al. 2020). Besides, with higher complexity comes a higher computational burden and the need for more data. To improve computation times in our case studies, we pre-calculated distances between patches for a set of resistance values rounded to one decimal, which saved us the need to compute distances between sites at each MCMC iteration. Regarding data requirement, occupancy models rely on detection/non-detection data that are not as informative on movements as GPS or telemetry data. Therefore, having enough variability in the studied process helps the model to converge. For instance, the delay between two seasons should be defined according to the dispersal speed of the focal species, to allow for dynamics in colonisation and extinction. We encounter convergence issues when the barrier is impermeable to dispersal (i.e., no individual are detected on the other side of a barrier). As landscape resistance increases, dispersal events get more difficult to detect. To cope with this issue, the surveys and seasons require to be defined to maximise detection of these rare dispersal events. We recommend building the sampling design so that the sampling effort is allocated homogeneously, on both sides of a barrier for example. Overall, our modelling framework offers a straightforward approach to efficiently assess connectivity. First, spatial occupancy models provide a data-driven evaluation of connectivity, based on detection/non-detection data which are readily available. Second, the Bayesian approach with MCMC explicitly propagates uncertainty, which provides reliable and robust inference on connectivity. Last, using R and especially NIMBLE ensures reproducibility of our approach (Riordan-Short, Pither, et Pither 2023). By providing a better understanding of population occurrence dynamics over space and time in fragmented landscapes, spatial occupancy models has become a valuable tool for species conservation and management.

## Conflict of Interest Statement

The authors declare no conflicts of interest.

## Appendix S1

**Bringing circuit theory into spatial occupancy models to assess landscape connectivity**, Maëlis Kervellec, Thibaut Couturier, Sarah Bauduin, Delphine Chenesseau, Pierre Defos du Rau, Nolwenn Drouet-Hoguet, Christophe Duchamp, Julien Steinmetz, Jean-Michel Vandel, Olivier Gimenez

**Figure S1.**
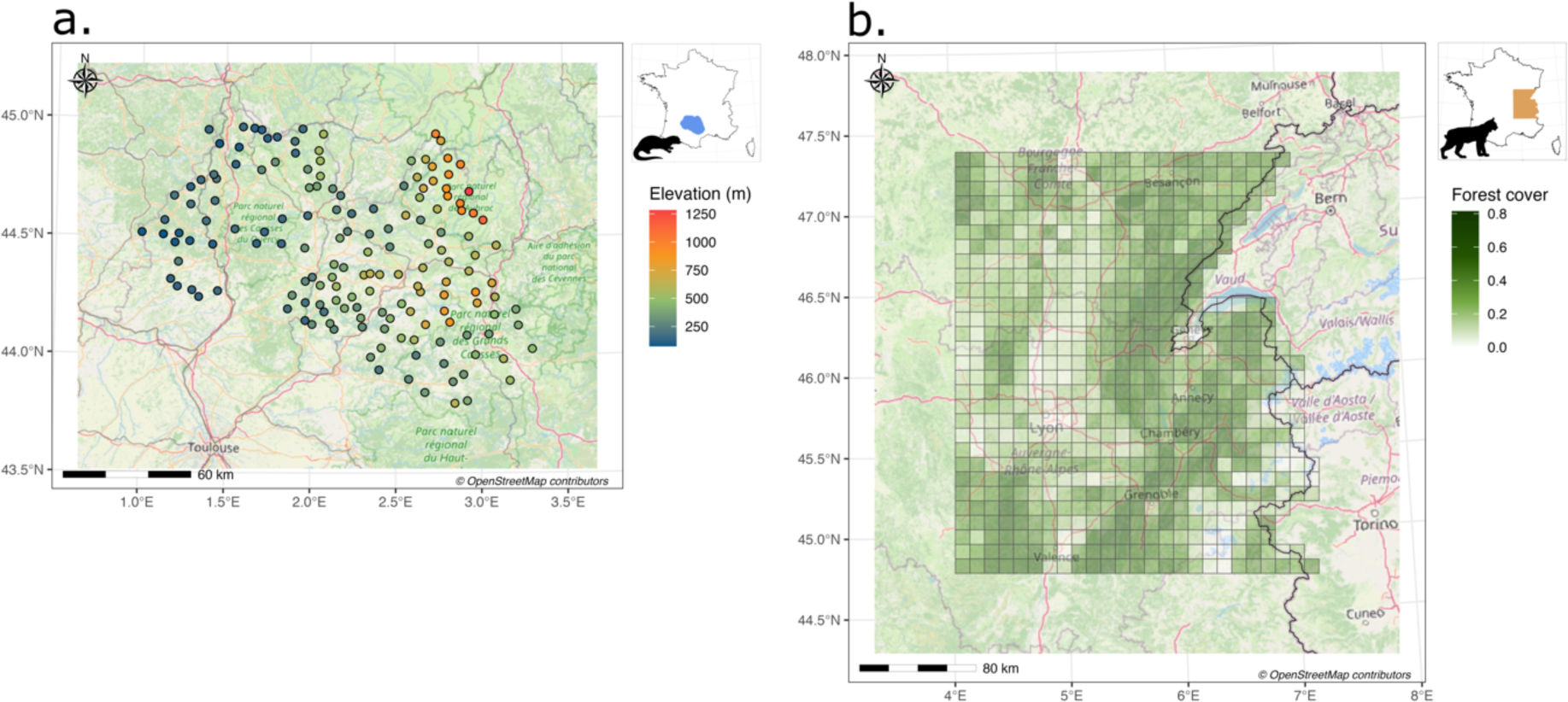
– Distribution of sampling sites for the 2 case studies in France. (a) 300m transect monitoring otter scats and tracks in the Massif Central coloured according to elevation (m). (b) 100km^2^ cells monitoring lynx presence in the Jura mountains and coloured according to the proportion of forest cover.

**Figure S2.**
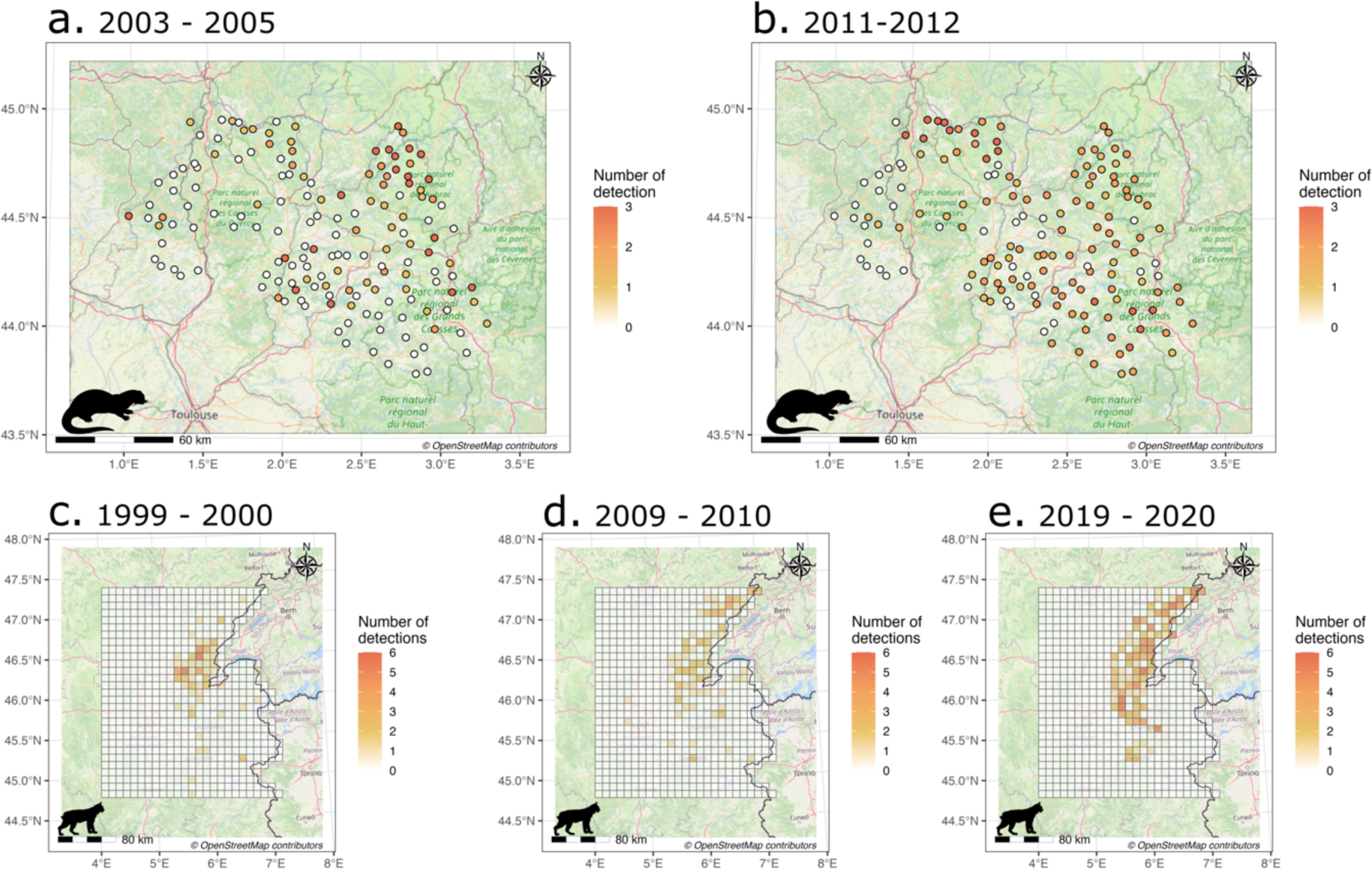
– Maps displaying the number of detections per season at each sampling site. The first row is for the otter in the Massif Central in 2003-2005 (a) and 2011-2012 (b). The second row is for lynx in the Jura mountains in 1999-2000 (c), 2009-2010 (d) and 2019-2020 (e).

**Figure S3.**
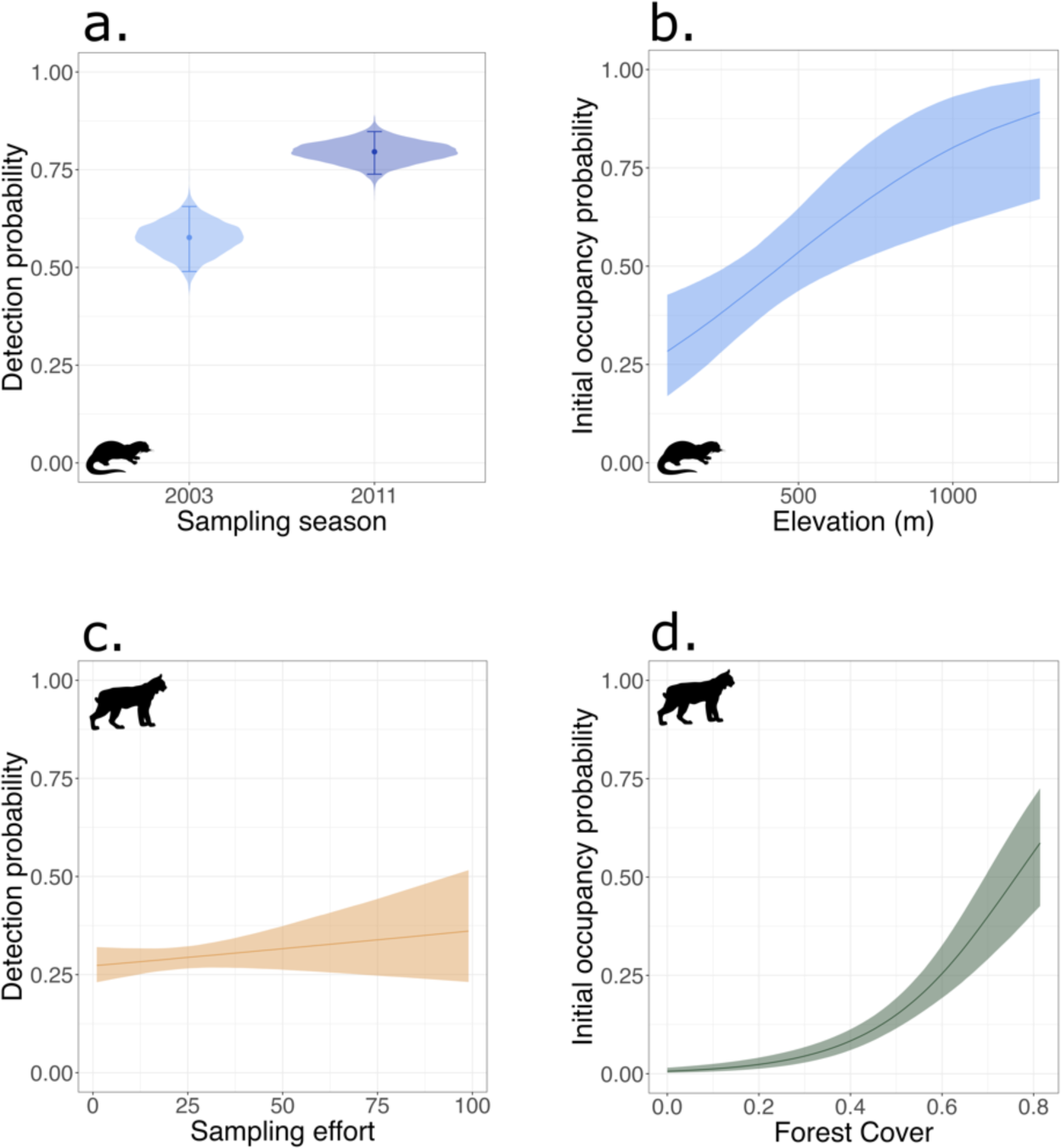
– Estimated relationships between detection, initial occupancy probabilities and covariates. First row: Otter detection probability as a function of sampling season (a), and initial ccupancy probability as a function of elevation (m) (b). Second row: lynx detection probability as function of sampling effort (c), and initial occupancy probability as a function of the forest cover d). Error bars and shaded areas are for 95% credible intervals. Points and lines represent posterior median.

**Figure S4.**
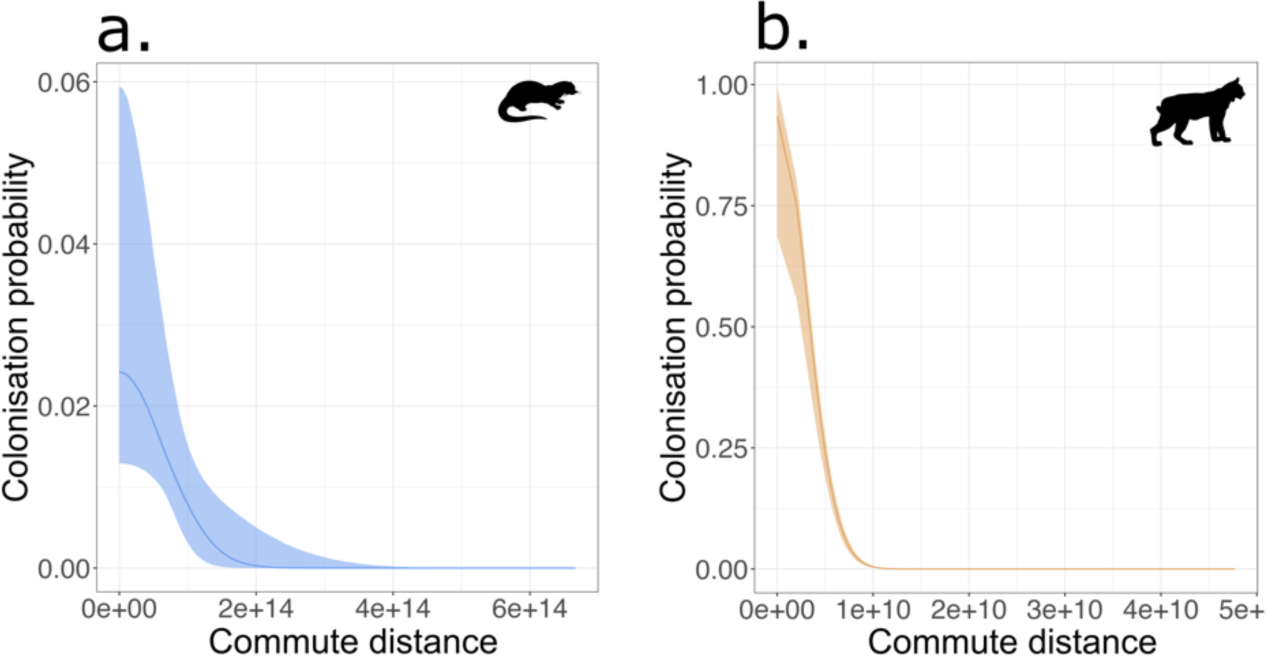
– Pairwise colonization probability (i.e. probability to be colonized by a given occupied site) as a function of commute distance computed from circuit theory with resistance parameter of respectively 3.9 and −0.2 for otter (a) and lynx (b). The lines represent posterior medians and shaded areas are for 95% credible intervals.

## Appendix S2

We present in the following document the code to implement dynamic occupancy models using NIMBLE (de Valpine et al., 2017). Reproducible examples with simulated data are available on Zenodo in the SpatialOccupancyModels folder.

DOI: 10.5281/zenodo.8376577

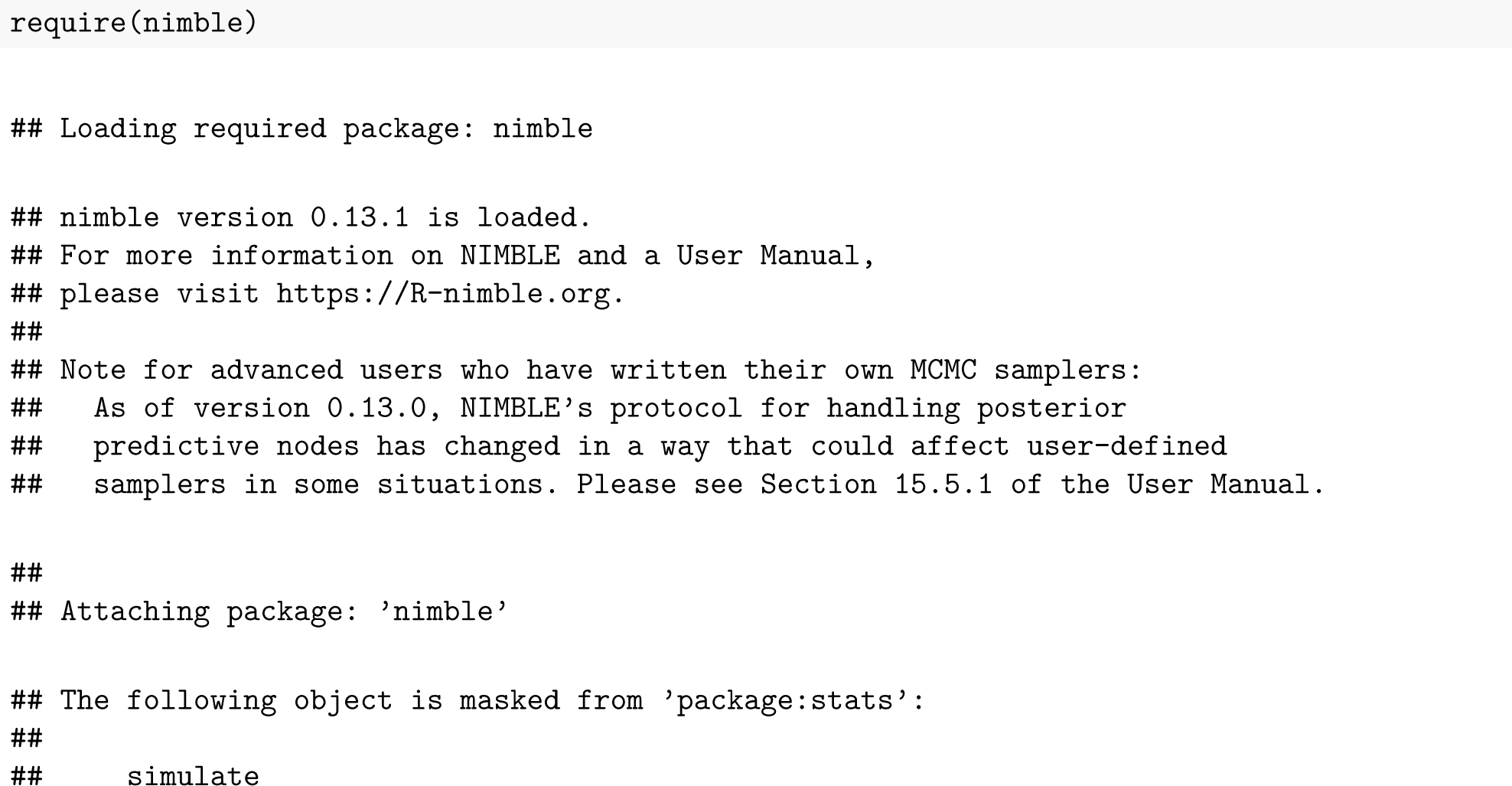

### 1. Dynamic occupancy model

You can find here the nimble code to run standard dynamic occupancy models. For more information on the model see Mackenzie et al. (2017).

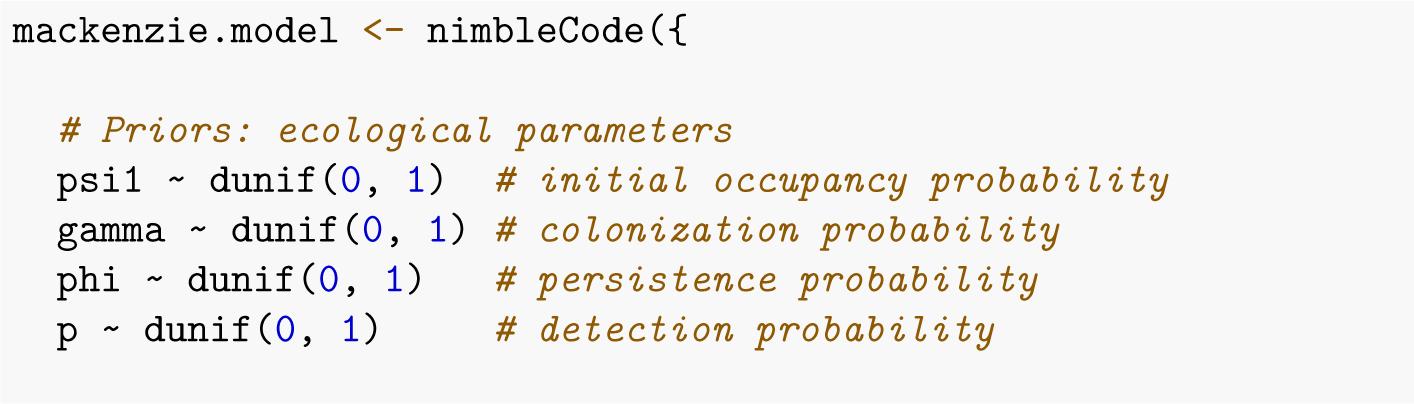
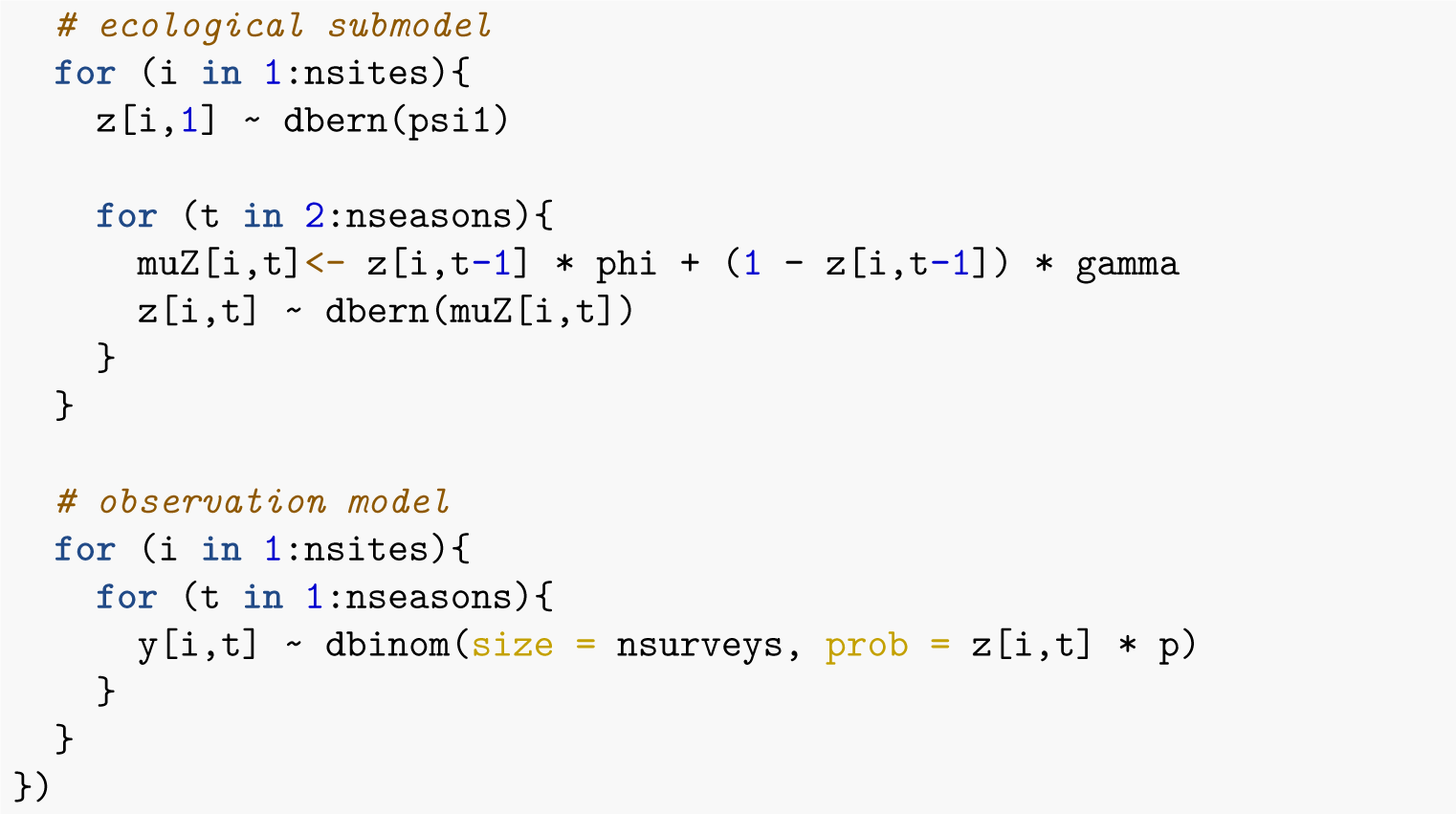

### 2. Spatial dynamic occupancy model

You can find here the nimble code to run spatial dynamic occupancy models using euclidean distance. For more information on the model see Chandler et al. (2015).

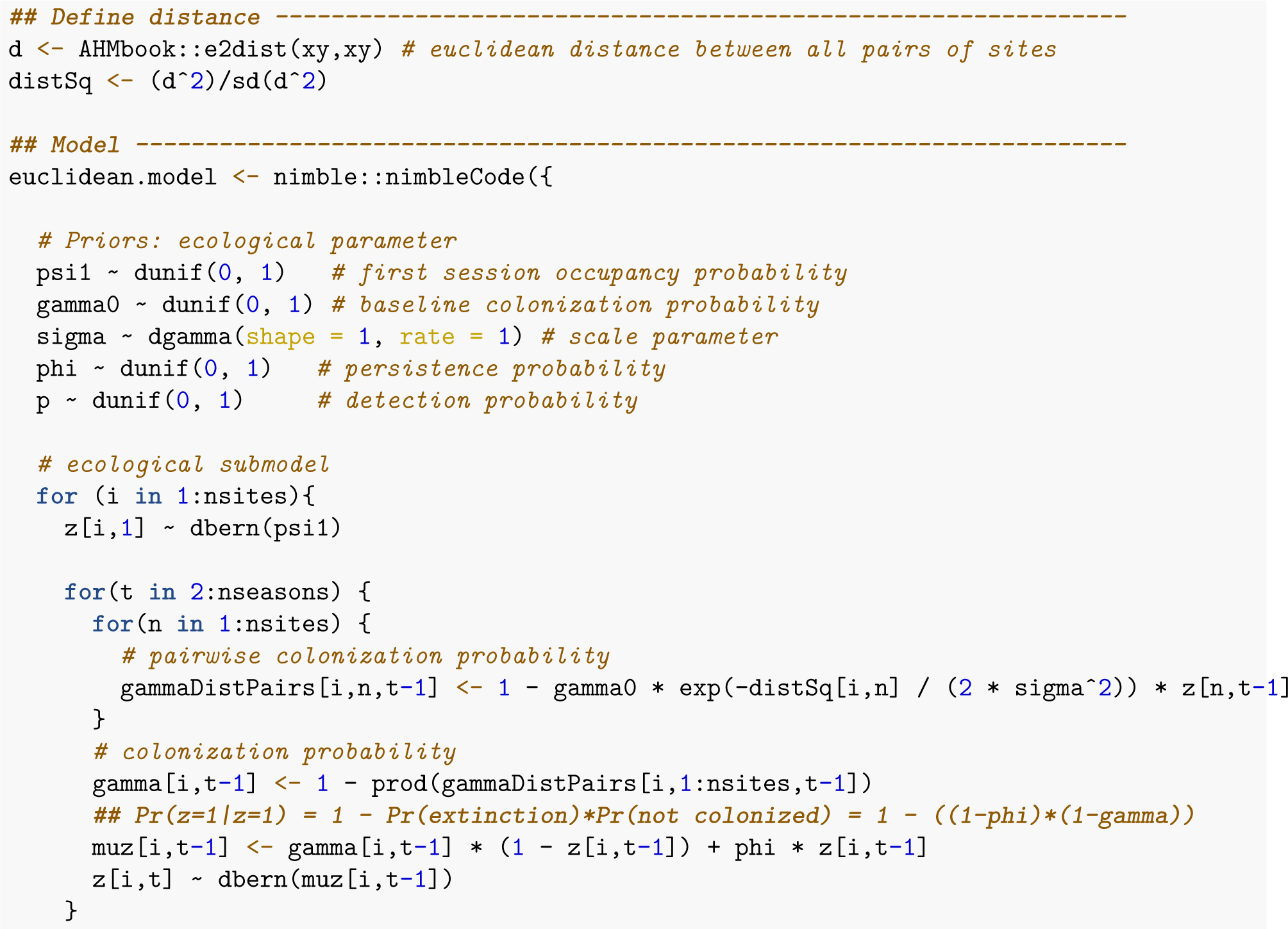
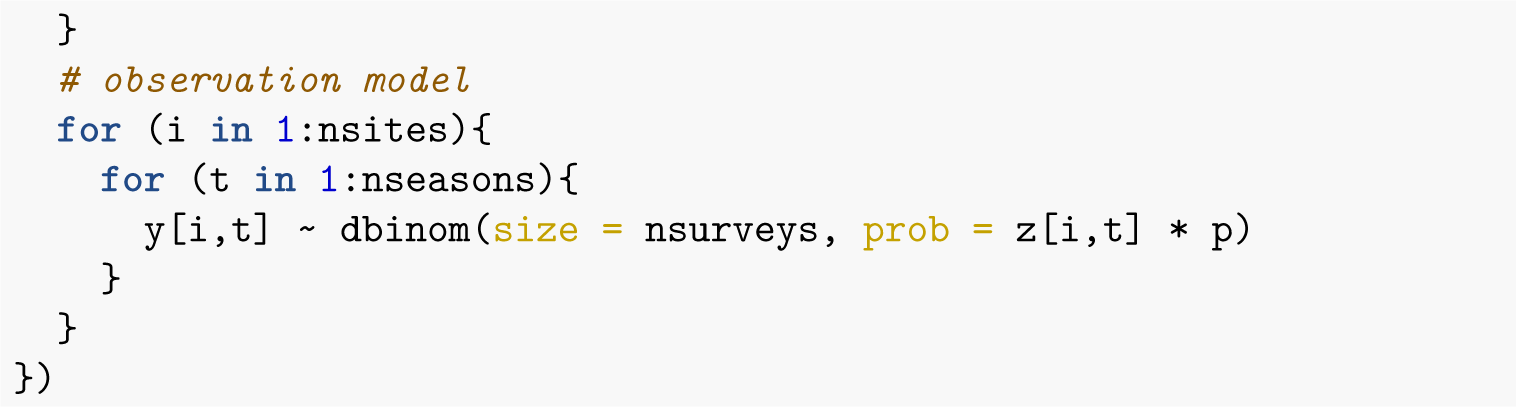

### 3. Spatial dynamic occupancy model accomodating Least Cost Path distance

You can find here the nimble code to run spatial dynamic occupancy models using least cost path distance. For more information on the model see Howell et al. (2018).

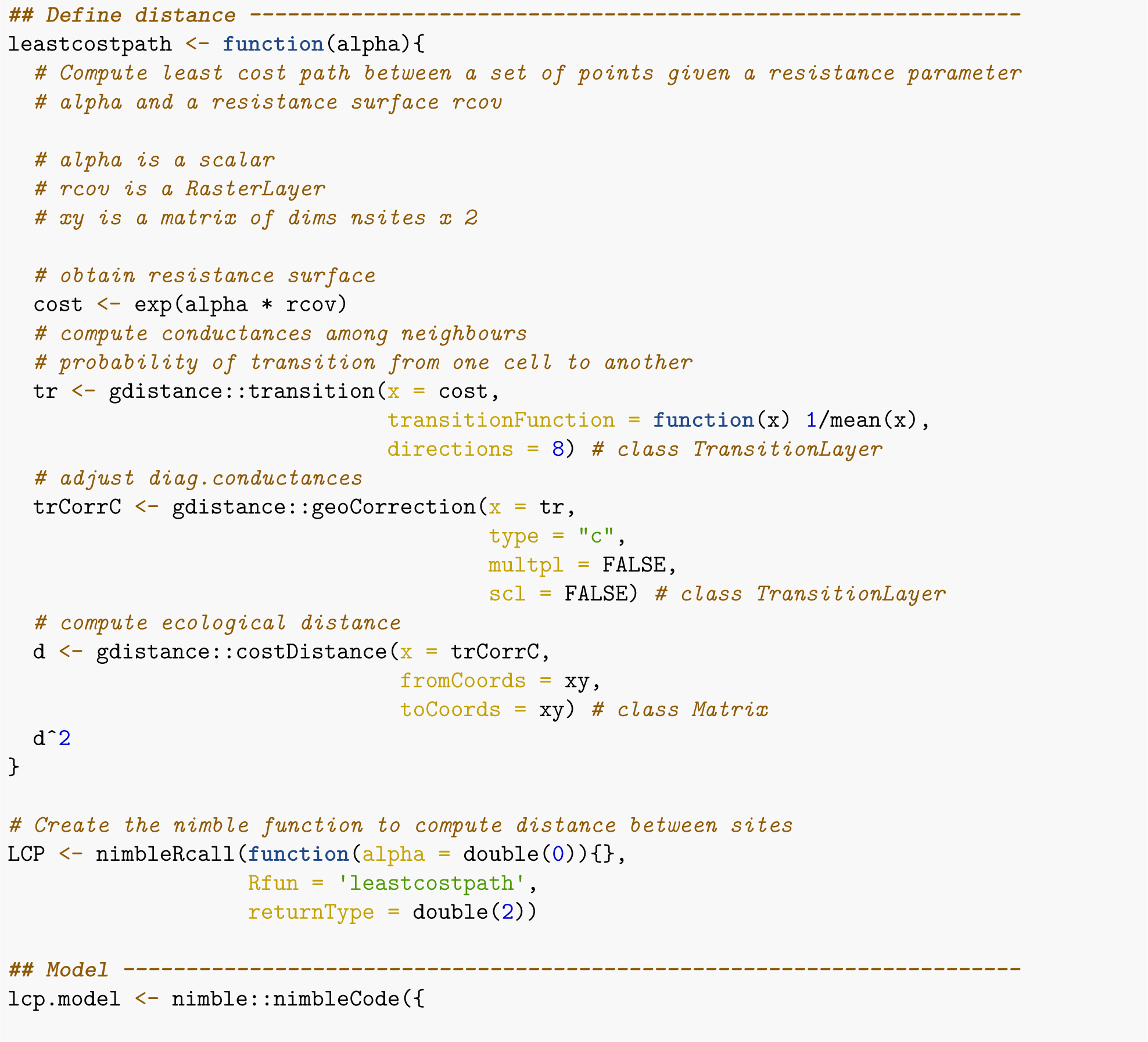
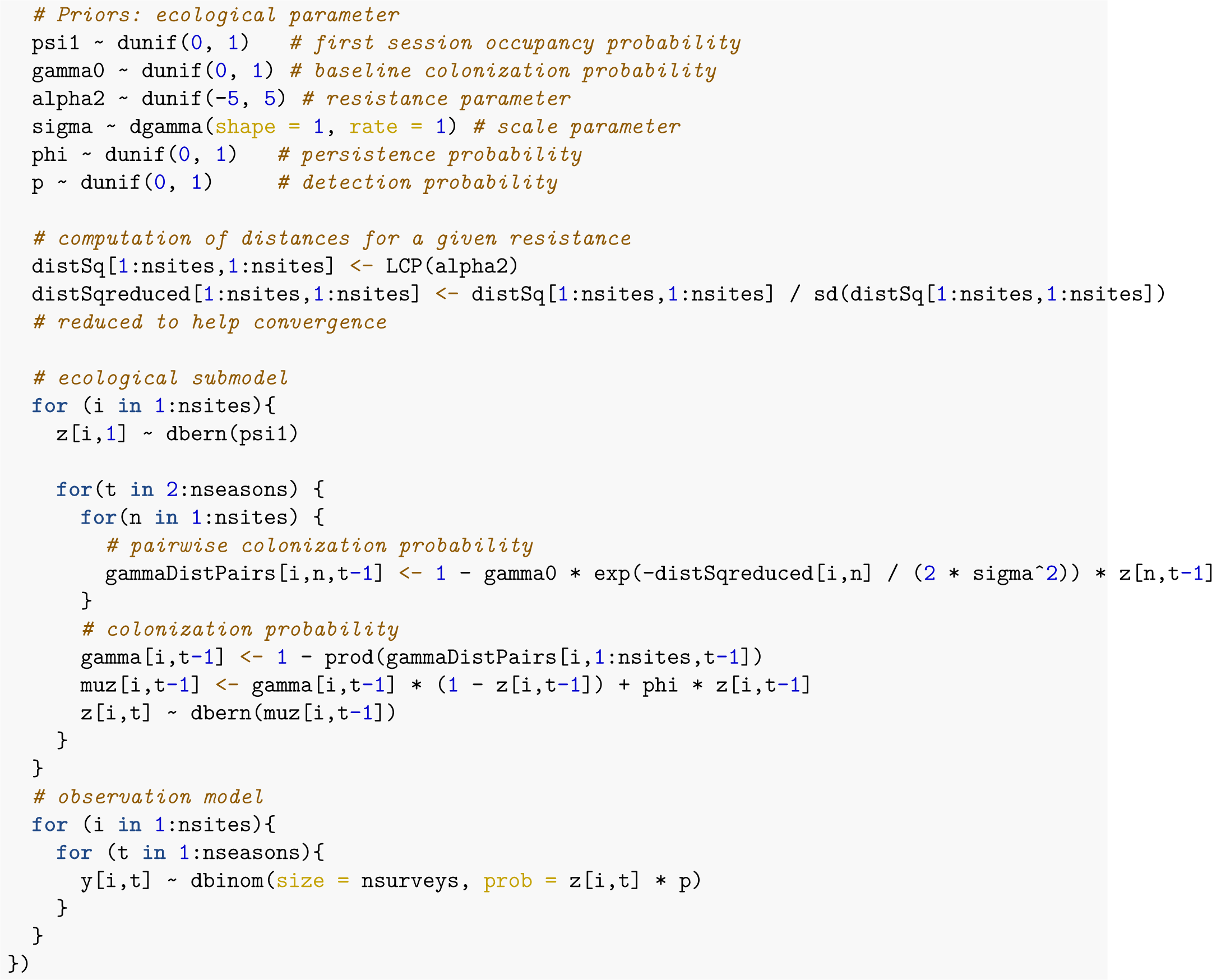

### 4. Spatial dynamic occupancy model accomodated with commute distance

You can find here the nimble code to run spatial dynamic occupancy models using commute distance from circuit theory. More information on the model are presented in the paper.

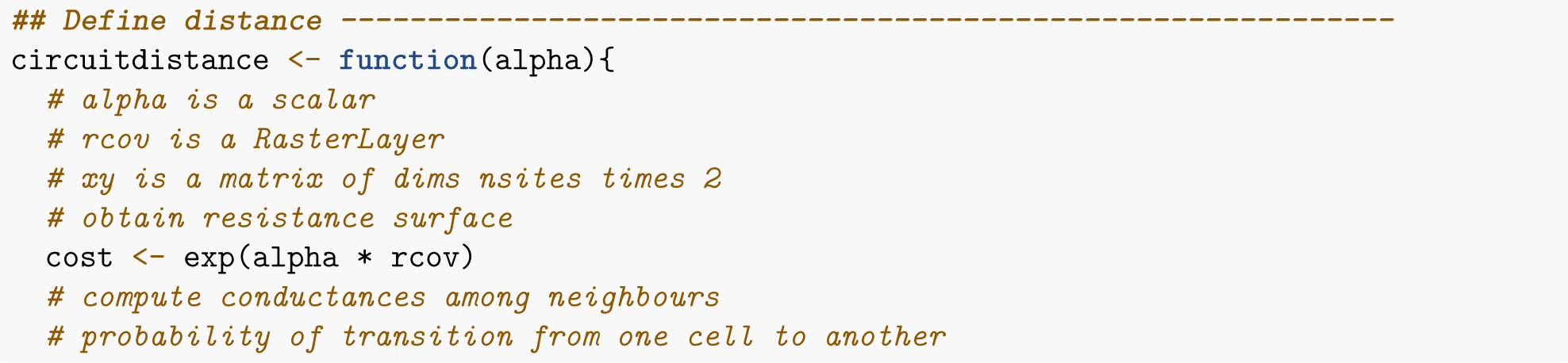
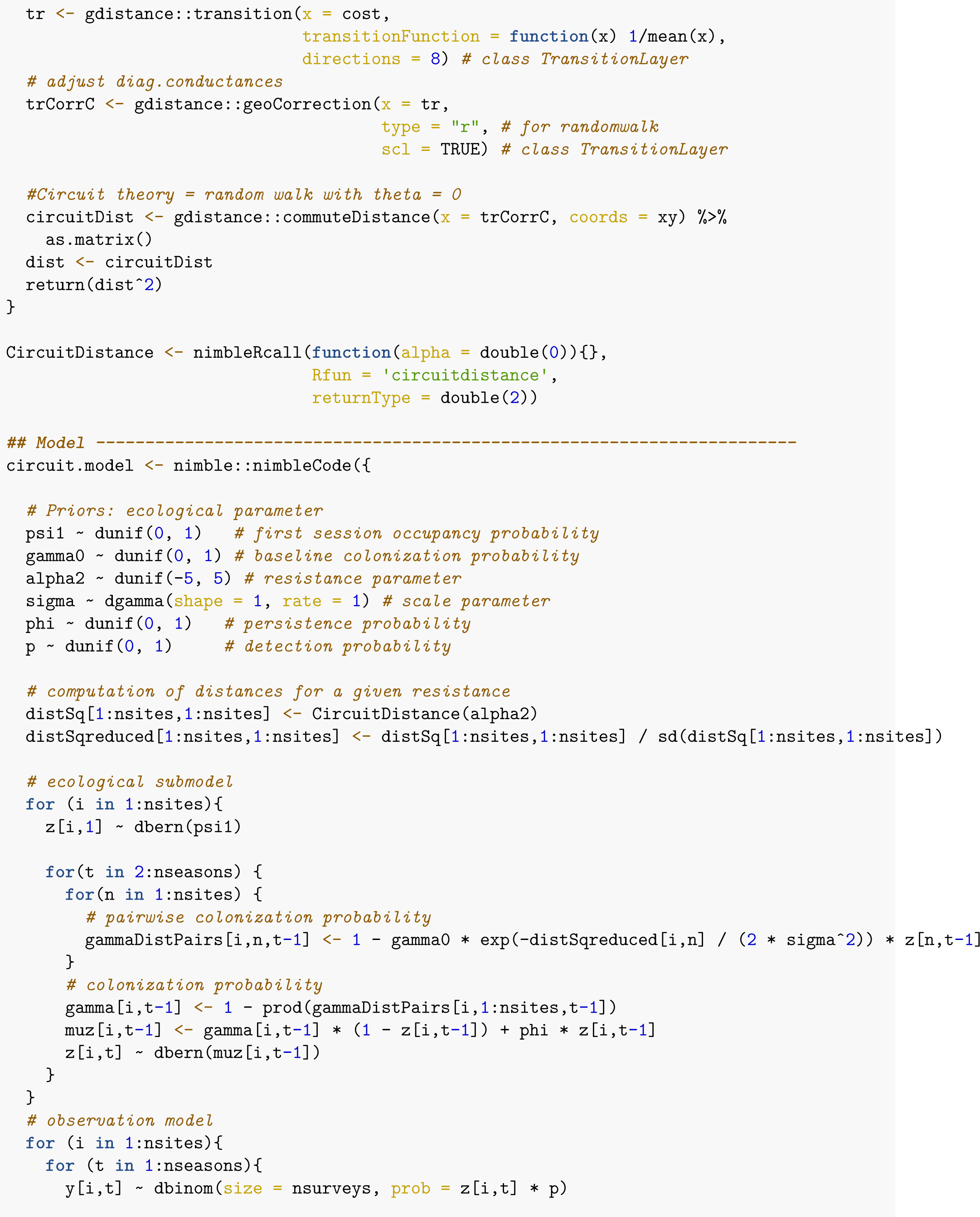
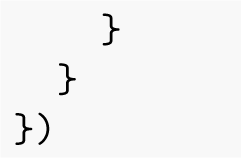

## Appendix S3

Trace plots and posterior density for spatial occupancy model applied to the otter data.

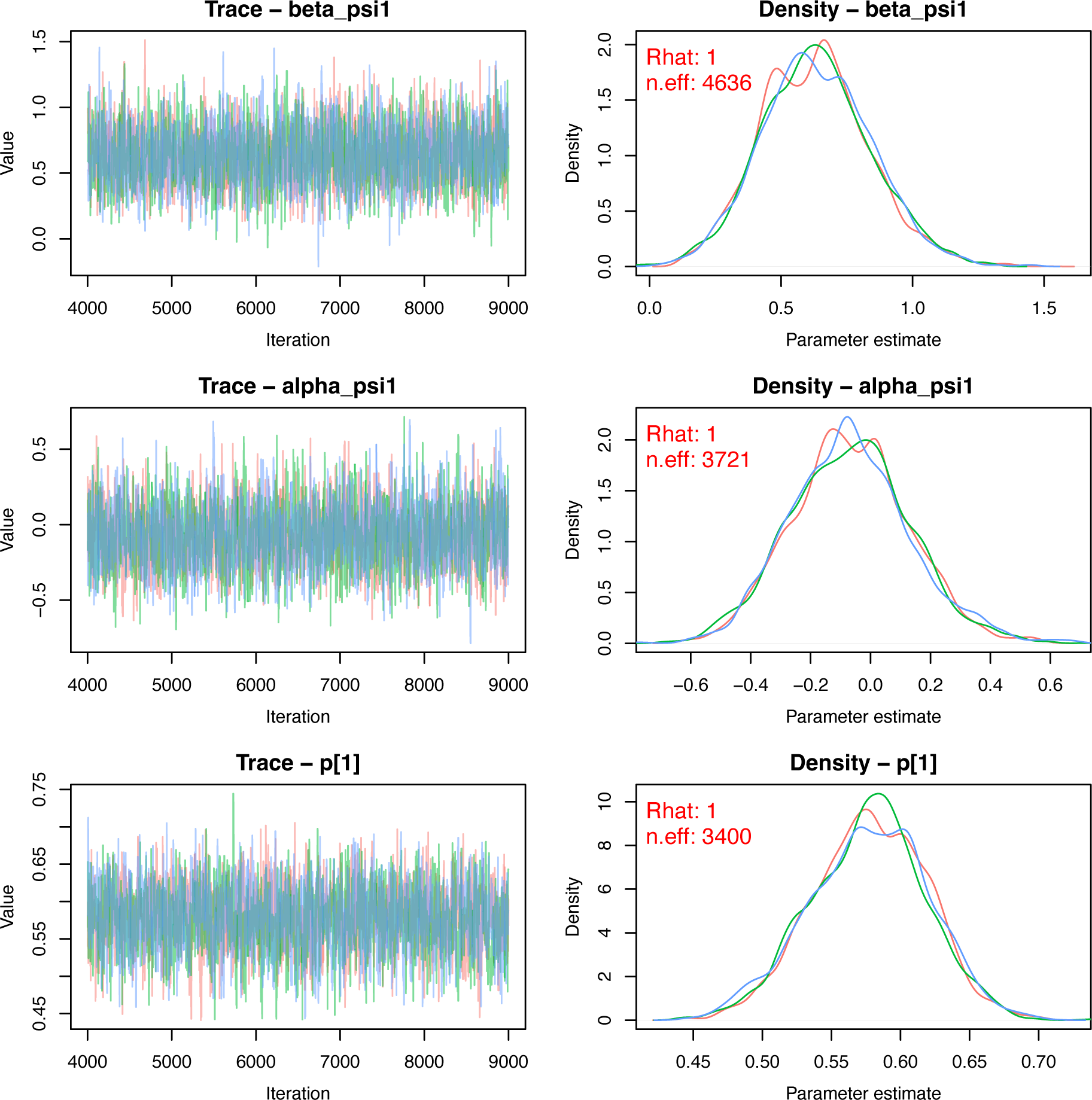
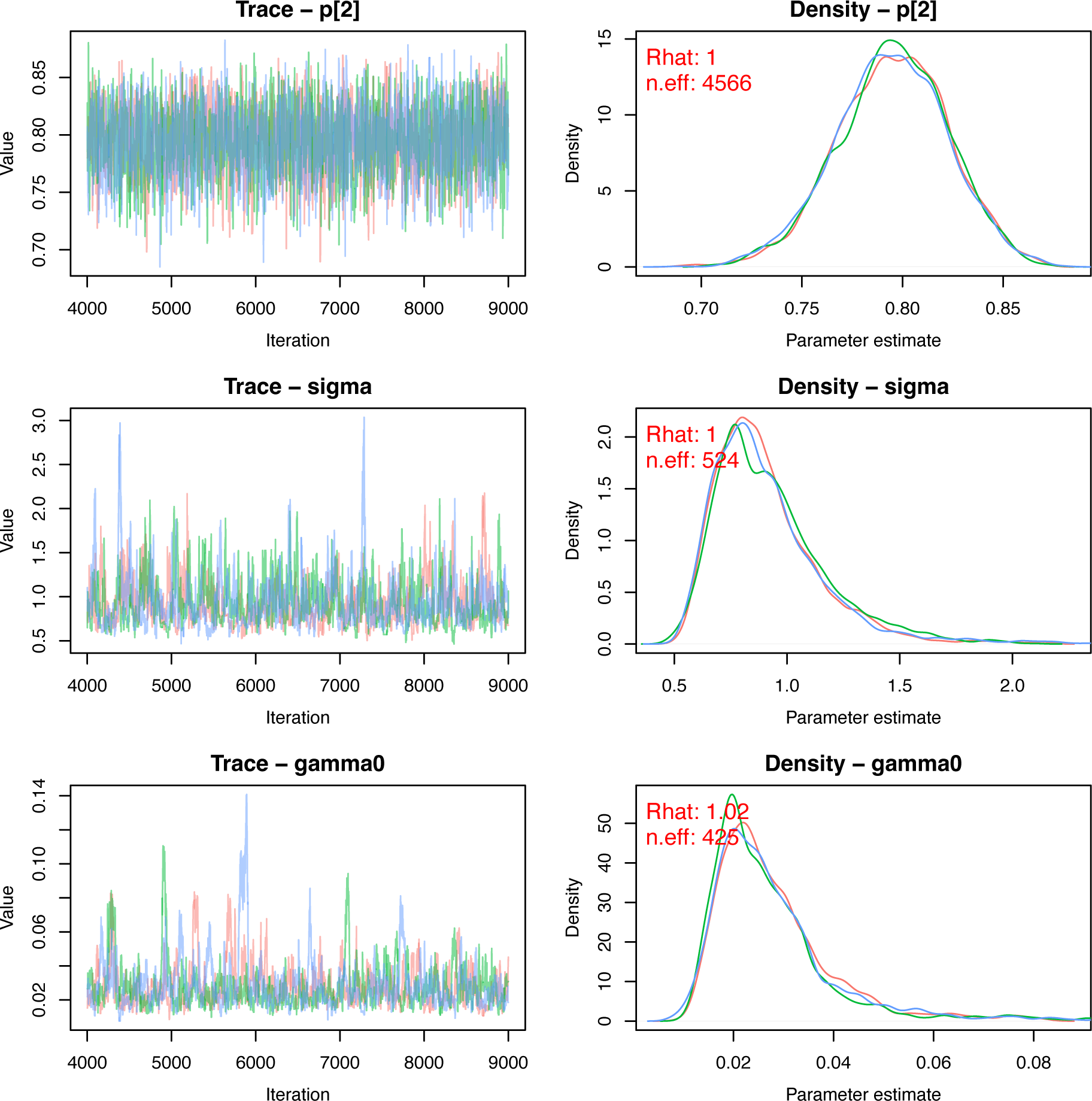
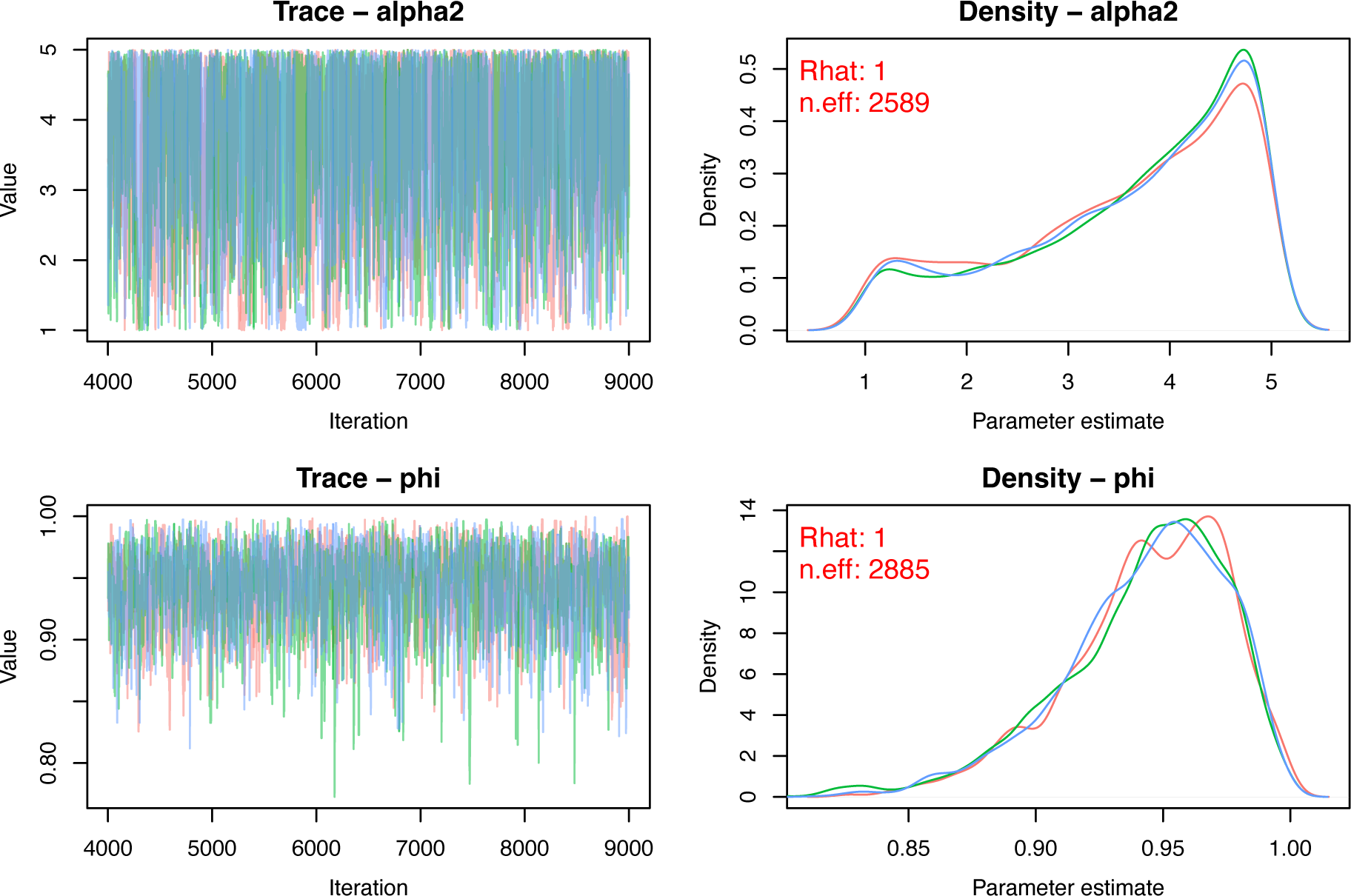

## Appendix S4

Trace plots and posterior density for spatial occupancy model applied to the lynx data.

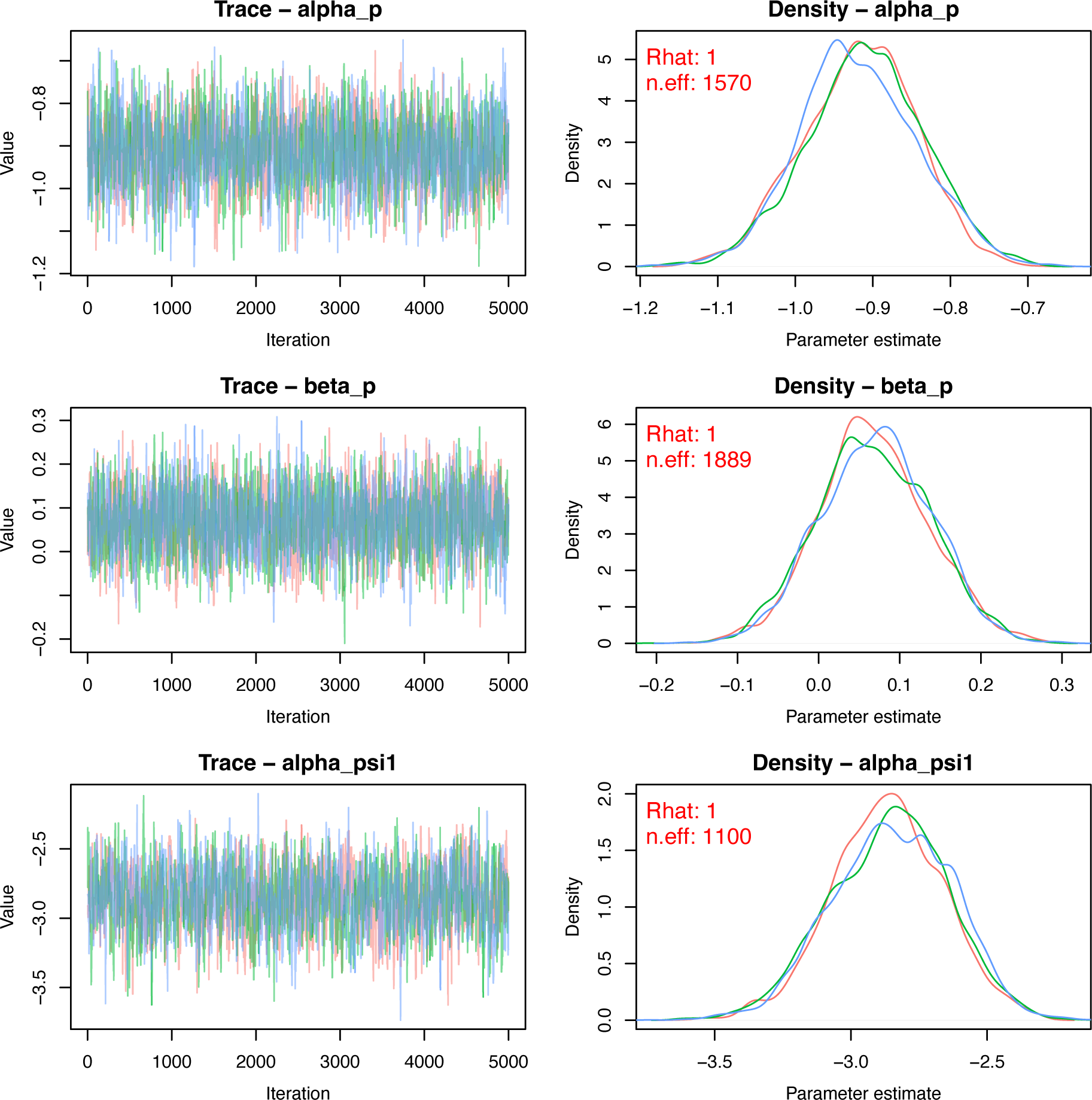
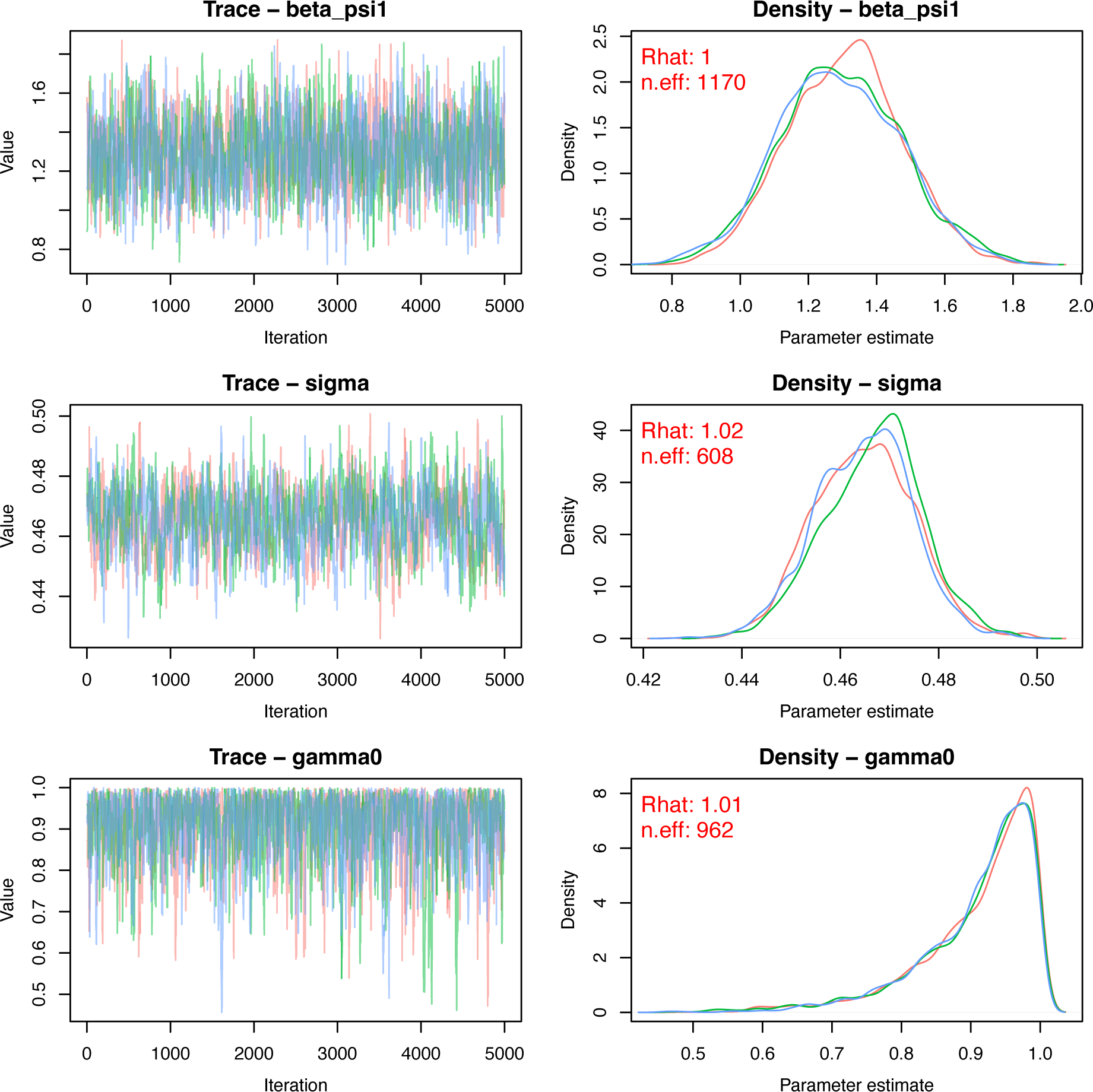
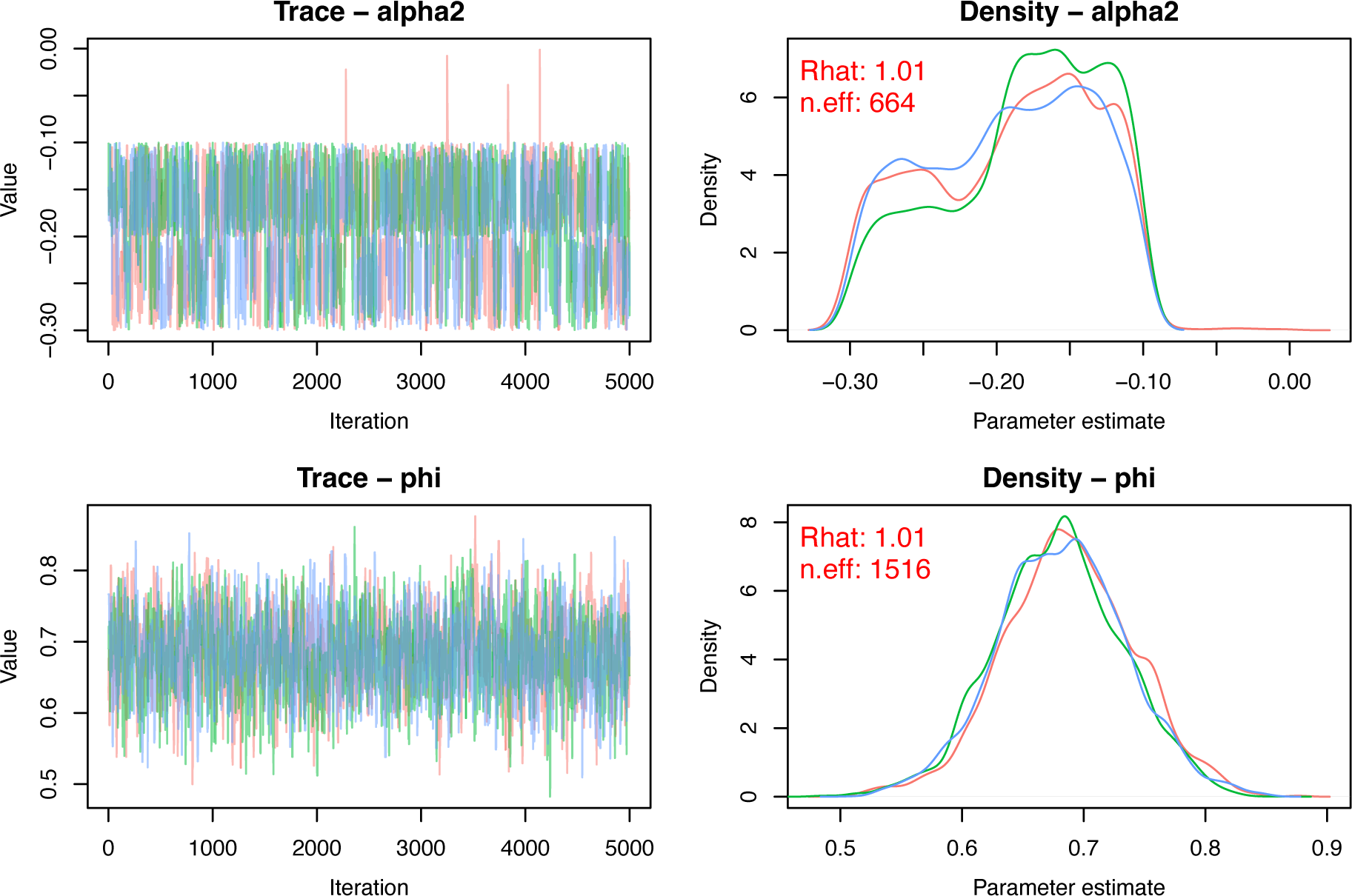

## Notes

https://zenodo.org/record/8376577

### Competing Interest Statement

The authors have declared no competing interest.

